# Biophysical characterization and modeling of *SCN1A* gain-of-function predicts interneuron hyperexcitability and a predisposition to network instability through homeostatic plasticity

**DOI:** 10.1101/2023.02.20.529310

**Authors:** Géza Berecki, Alexander Bryson, Tilman Polster, Steven Petrou

**Affiliations:** Ion Channels and Disease Group, The Florey Institute of Neuroscience and Mental Health, The University of Melbourne, Parkville, VIC 3052, Australia; Department of Neurology, Austin Health, Heidelberg, VIC 3084, Australia; Krankenhaus Mara, Bethel Epilepsy Centre, Department of Epileptology, Medical School, Bielefeld University, Campus Bielefeld-Bethel, Bielefeld, Germany; Praxis Precision Medicines, Inc., Cambridge, MA 02142, USA; Department of the Florey Institute, University of Melbourne, Parkville, VIC 3050, Australia

**Keywords:** *SCN1A*, early-onset developmental and epileptic encephalopathy, dynamic action potential clamp, spiking cortical network model

## Abstract

*SCN1A* gain-of-function variants are associated with early onset developmental and epileptic encephalopathies (DEEs) that possess distinct clinical features compared to Dravet syndrome caused by *SCN1A* loss-of-function. However, it is unclear how *SCN1A* gain-of-function may predispose to cortical hyper-excitability and seizures. Here, we first report the clinical features of a patient carrying a *de novo SCN1A* variant (T162I) associated with neonatal-onset DEE, and then characterize the biophysical properties of T162I and three other *SCN1A* variants associated with neonatal-onset or early infantile DEE (I236V, P1345S, R1636Q). In voltage clamp experiments, three variants (T162I, P1345S and R1636Q) exhibited changes in activation and inactivation properties that enhanced window current, consistent with gain-of-function. Dynamic action potential clamp experiments utilising model neurons incorporating Na_v_1.1. channels supported a gain-of-function mechanism for all four variants. Here, the T162I, I236V, P1345S, and R1636Q variants exhibited higher peak firing rates relative to wild type and the T162I and R1636Q variants produced a hyperpolarized threshold and reduced neuronal rheobase. To explore the impact of these variants upon cortical excitability, we used a spiking network model containing an excitatory pyramidal cell (PC) and parvalbumin positive (PV) interneuron population. *SCN1A* gain-of-function was modeled by enhancing the excitability of PV interneurons and then incorporating three simple forms of homeostatic plasticity that restored pyramidal cell firing rates. We found that homeostatic plasticity mechanisms exerted differential impact upon network function, with changes to PV-to-PC and PC-to-PC synaptic strength predisposing to network instability. Overall, our findings support a role for *SCN1A* gain-of-function and inhibitory interneuron hyperexcitability in early onset DEE. We propose a mechanism through which homeostatic plasticity pathways can predispose to pathological excitatory activity and contribute to phenotypic variability in *SCN1A* disorders.

## INTRODUCTION

The *SCN1A* gene encodes the voltage-gated sodium channel Na_v_1.1 which is preferentially expressed in GABAergic interneurons and plays a critical role in regulating cortical excitability (Duflocq et al., 2008; Lorincz and Nusser, 2008; Ogiwara et al., 2007). Loss-of-function *SCN1A* mutations are associated with epilepsy syndromes of varying clinical severity, from the mild genetic epilepsy with febrile seizures plus (GEFS+) (Myers et al., 2018) to the severe developmental and epileptic encephalopathy (DEE) Dravet syndrome (Connolly, 2016; Mulley et al., 2005). Although a clear relationship between the severity of the clinical phenotype and Na_v_1.1 channel dysfunction is not uniformly observed, epilepsy syndromes associated with *SCN1A* loss-of-function are thought to arise through network hyper-excitability due to disinhibition of excitatory pyramidal cells, in part due to impaired parvalbumin (PV)-positive interneuron function (Ragsdale, 2008). Nav1.1 channels are also expressed in GABAergic somatostatin (SST)-positive and vasoactive intestinal peptide (VIP)-positive interneurons, and Na_v_1.1 channel dysfunction is now recognised to exert a synergistic impact on cortical excitability and epileptogenesis in Dravet Syndrome through dysfunction of multiple interneuron subtypes (Goff and Goldberg, 2019; Rubinstein et al., 2015; Tai et al., 2014). Interneurons are known to regulate the gain of pyramidal cells and tightly regulate network excitatory-inhibitory (EI) balance (Isaacson and Scanziani, 2011; Klausberger et al., 2003).

Interestingly, *SCN1A* gain-of-function variants also result in a disease spectrum ranging from familial hemiplegic migraine to severe forms of early onset DEE (Brunklaus et al., 2022; Mantegazza and Cestele, 2018). However, the DEE phenotype associated with *SCN1A* gain-of-function is distinct from that associated with *SCN1A* loss-of-function, as first described in nine unrelated children, eight of whom carried the T226M variant and one the P1345S variant (Sadleir et al., 2017). These cases were characterized by early infantile seizure onset, profound intellectual disability, and movement disorders. In a subsequent study, 20 additional *SCN1A* gain-of-function variants were identified and associated with early infantile DEE with or without movement disorder (EIDEE/MD) or severe neonatal DEE with movement disorder and arthrogryposis (NDEEMA) clinical phenotypes. In heterologous expression systems, the functional evaluation of *SCN1A* variants associated with NDEEMA (T162I, I236V) and EIDEE/MD (R1636Q, I1498T) revealed changes in biophysical characteristics relative to wild-type that predispose to gain-of-function. (Brunklaus et al., 2022; Clatot et al., 2022; Matricardi et al., 2023).

However, as demonstrated for the T226M mutant, enhanced Na_v_1.1 current may exert paradoxical effects upon neuronal firing leading to a transient increase of firing rates followed by enhanced susceptibility to depolarisation block (Berecki et al., 2019). Furthermore, it is unclear how these changes disrupt network function, and it is possible that homeostatic adaptations in response to intrinsic alterations of neuron function can produce unexpected effects upon cortical excitability. Indeed, homeostatic adaptations to cortical networks that act to restore normal firing rates may play a crucial role in epileptogenesis (Keck et al., 2017; Lignani et al., 2020; Nasrallah et al., 2022).

In this study, we characterized the biophysical channel properties and the impact upon neuronal excitability of four *SCN1A* variants associated with NDEEMA (T162I, I236V) and EIDEE/MD (P1345S, R1636Q). We show that all variants exhibit changes in biophysical properties relative to wild-type, consistent with gain-of-function leading to increased action potential firing of hybrid model PV interneurons in dynamic action potential clamp (DAPC) experiments. To explore the potential impact of these variants upon cortical activity, we first incorporated the changes to PV interneuron excitability associated with *SCN1A* gain-of-function within a spiking network model containing an excitatory pyramidal cell (PC) and a PV interneuron population. We then implemented three forms of homeostatic plasticity that act to restore PC firing rates: reductions of PV-to-PC synaptic strength (Model 1), increased PC-to-PC synaptic strength (Model 2), or increased intrinsic PC excitability through a reduction of firing rate threshold (Model 3). Although all models exhibited similar baseline firing rates, changes to PV-to-PC and PC-to-PC synaptic strength predisposed to excitation-inhibition imbalance and synchronous excitatory activity in response to external stimulation.

Overall, our findings support a pathogenic role for *SCN1A* gain-of-function and PV interneuron hyperexcitability in early onset DEE, and we propose a mechanism through which homeostatic plasticity may differentially predispose to cortical excitability. It is possible that selective modulation of these pathways may be of therapeutic benefit by driving network adaptations into a less excitable state.

## METHODS

### Identification of variants and clinical data

The study was conducted in accordance with the Declaration of Helsinki and approved by the Ethics Committee of the University of Leipzig, Germany (224/16-ek and 402/16-ek). Written informed consent for genetic testing and the publication of findings after advice and information about the study was obtained from the legal representatives of the patient involved in the study. Written informed consent was obtained for all individuals whose previously unpublished clinical data is presented here. Clinical information for the individual carrying the recurrent T162I mutation was obtained from clinical visits, hospital admissions and medical records by a paediatric epileptologist (TP). Clinical data of patients with T162I, I236V, P1345S, and R1636Q variants had been previously reported (Brunklaus et al., 2022; Clatot et al., 2022; Matricardi et al., 2023; Sadleir et al., 2017).

### *SCN1A* variants and site-directed mutagenesis

The human wild-type *SCN1A* cDNA used in this study (NCBI Ref. Seq. NM_001165963.2), a kind gift from Dr. AL George Jr. (Department of Pharmacology, Northwestern University). The *SCN1A* sequence contains silent mutations to increase DNA stability for bacterial expression without affecting the protein sequence, as previously described (Berecki et al., 2019). QuikChange Lightning Site-Directed Mutagenesis Kit (Agilent Technologies, Santa Clara, CA) was performed to introduce the mutations into the wild-type *SCN1A*, using custom made primers (Bioneer Pacific Australia). Sequences were as follows: for c.485C>T, p.T162I, 5’-GGACAAAGAATGTAGAATACACCTTCA**T**AGGAATATATACTTTTG-3’ (forward, F) and 5’-CAAAAGTATATATTCCT**A**TGAAGGTGTATTCTACATTCTTTGTCC-3’ (reverse, R); for c.706A>G, p.I236V, 5’-CAGGCCTGAAAACC**G**TTGTGGGAGCCCTG-3’ (F) and 5’-CAGGGCTCCCACAA**C**GGTTTTCAGGCCTG-3’ (R); for c.4033C>T, p.P1345S, 5’-TAAATGCTCTTTTAGGAGCCATT**T**CATCTATCATGAATGTACTTCT-3’ (F) and 5’-AGAAGTACATTCATGATAGATG**A**AATGGCTCCTAAAAGAGCATTTA-3’ (R); for c.4907G>A, p.R1636Q, 5’-CCCCTACCCTGTTCC**A**AGTGATCCGTCTTGC-3’ (F) and 5’-GCAAGACGGATCACT**T**GGAACAGGGTAGGGG-3’ (R); mutated bases are in bold. The full sequences of all *SCN1A* variants were validated by automated DNA sequencing (Australian Genome Research Facility, Melbourne).

### Cell culture and transient transfection

Chinese hamster ovary (CHO) cells were maintained in Dulbecco’s Modified Eagle Medium: Nutrient Mixture F-12 (Thermo Fisher Scientific, Waltham, MA) supplemented with 10% (v/v) fetal bovine serum (Thermo Fisher Scientific) and 50 IU/ml penicillin (Thermo Fisher Scientific) at 37°C and 5% CO_2_. Cells were co-transfected at ∼80% confluence with wild-type (4 μg) or mutant *SCN1A* construct (4 μg) and enhanced green fluorescent protein (eGFP (1 μg) using Lipofectamine 3000 Reagent (Thermo Fisher Scientific) and incubated under the culture conditions above for 24 hours. Transfected cells were treated with TrypLE Express Reagent (Thermo Fisher Scientific), plated on 13 mm diameter glass coverslips (Menzel-Gläser, Thermo Fisher Scientific), and incubated at 30°C in 5% CO_2_.

### Electrophysiology

Sodium currents were recorded in the whole-cell patch clamp configuration, 48 to 72 hours after transfection, using an Axopatch 200B amplifier (Molecular Devices, Sunnyvale, CA) controlled by a pCLAMP 9/DigiData 1440 acquisition system (Molecular Devices). The extracellular bath solution contained 145 mM NaCl, 5 mM CsCl, 2 mM CaCl_2_, 1 mM MgCl_2_, 5 mM glucose, 5 mM sucrose, 10 mM Hepes (pH = 7.4 with NaOH), whereas the intracellular solution contained 5 mM CsCl, 120 mM CsF, 10 mM NaCl, 11 mM EGTA, 1 mM CaCl_2_, 1 mM MgCl_2_, 2 mM Na_2_ATP, 10 mM Hepes (pH = 7.3 with CsOH). Fire-polished borosilicate patch pipettes (GC150TF-7.5, Harvard Apparatus Ltd.) typically exhibiting resistance values of 1.1−1.5 MΩ were used, and series resistances were compensated 85 to 90% in all cases. Membrane currents and potentials were low-pass filtered at 10 kHz and sampled at 50 kHz. In Leak and capacitive currents were subtracted using a −P/4 pulse protocol, unless mentioned otherwise. The voltage clamp protocols are shown in the figures and explained in the Results section. To minimize possible voltage errors, CHO cells expressing peak sodium current amplitudes smaller than 2 nA or larger than 10 nA were excluded from biophysical characterization datasets as previously described (Berecki et al., 2018). The current-voltage (I−V) relationships, the parameters of the voltage dependence of (in)activation, persistent current (I_NaP_), and the time constant of recovery from fast inactivation were determined as previously described (Berecki et al., 2018).

Dynamic action potential clamp experiments were performed by implementing scaled sodium currents of individual Na_v_1.1 channel variants in a biophysically realistic AIS model as previously described (Berecki et al., 2019). The in-silico sodium conductance of the AIS model was set to zero (gNa_v_1.6 = 0), whereas the in silico K_v_ channel conductance was set to default (gK_v_ = 1) or gK_v_ = 2. Action potential firing of the AIS hybrid model neuron was elicited from a holding membrane potential value of −70 mV using step current injections in 2-pA increments. The threshold was estimated using Axograph X (Axograph Scientific, Sydney, Australia), as the point on the action potential rising phase where the first derivative (dV/dt) of the voltage trajectory reached 20 mV/ms. The firing frequency, rheobase, action potential width and decay time were determined using the Clampfit module of pCLAMP 10, as described previously (Berecki et al., 2018).

### Spiking and simplified network models

The spiking network model was adopted from Bryson et al (Bryson et al., 2021) and is available at https://osf.io/75vky/. In brief, excitatory pyramidal cell (PC) models contained a somatic and dendritic compartment, five voltage-dependent mechanisms described with the Hodgkin-Huxley formulation and were optimised to exhibit realistic electrophysiological properties in response to constant-current stimuli (Druckmann et al., 2011). PV interneurons were modeled using the formulation developed by Izhikevich at al. and synapses with a bi-exponential formalism (Izhikevich, 2003). The network model contained 1800 excitatory PC neurons and 450 (20%) inhibitory interneurons. PC-to-PC connection probability was set to 15% and recurrent connectivity between the PC and interneuron population to 25%. Synaptic weights possessed a log-normal distribution and means and variances of synaptic weights and latencies were identical to previously published values (Bryson et al., 2021). The network was stimulated with Poisson-distributed excitatory synaptic inputs that elicited a mean excitatory and inhibitory firing rate of 0.3 Hz and 2 Hz, respectively, comparable to firing rates observed in-vivo (Barth and Poulet, 2012). Simulations were performed using NetPyne using a fixed integration time step of 0.025 ms (Dura-Bernal et al., 2019). External stimulation was applied to all PC neurons with a 1.5 nA current stimulus of 10 ms duration. Interneuron excitability was enhanced to model *SCN1A* GoF by applying a constant current input that produced a 25% reduction of firing threshold. Homeostatic plasticity was modelled by either reducing PV-to-PC synaptic strength (Model 1), increasing PC-to-PC synaptic strength (Model 2), or reducing PC rheobase by applying a constant current stimulus (Model 3) to restore a baseline population excitatory firing rate of ∼0.3 Hz. Peak feedback inhibition, recurrent excitation and EI ratio were measured by recording currents generated by all inhibitory and excitatory synapses onto all PC neurons and averaged across 20 simulations initiated with a different random seed. The sensitivity analysis was performed using a grid search by varying each parameter (PV-to-PC synaptic strength, PC-to-PC synaptic strength and PC rheobase) by 25% increments to the value that restored PC firing rates. This led to 5 × 5 × 5 = 125 models in total. Network stability (Fig. 5c) was assessed by measuring the number of PC spikes within a 10 ms window after external stimulation was applied to each model, averaged over 5 simulations initiated with a different random seed.

The rate-based network was described using a Wilson-Cowan model with one excitatory (E) and inhibitory (I) population, linear input-output relationships (gain equal to 1) and constant external excitatory input. All synaptic weights were set to 1.0 except E-to-E (*W*_*EE*_) and I-to-E (*W*_*EI*_) synapses which were set to 1.25 and 1.75, respectively, and time constants of each population were set to 20 ms. External input was applied to reproduce firing rates identical to the mean rates of the spiking network. Homeostatic plasticity was modelled by either increasing *W*_*EE*_, reducing *W*_*EI*_, or reducing the mean E firing threshold. Network stability in Fig. 6d was calculated by computing the real part of the eigenvalue of highest value associated with the model’s Jacobian matrix for values of *W*_*EE*_ between 0.75 and 1.75 and *W*_*EI*_ between 0.2 to 1.4. External stimulation to the E population comprised a 10 nA stimulus of 1 ms duration. Simulations were performed with XPPAUT (Ermentrout and Mahajan, 2003).

### Statistical analysis

Voltage clamp and DAPC data were plotted in Origin 2022 (Microcal Software Inc., Northampton, MA). Mean and standard error of the mean (SEM) were used to describe the variability within groups; n values show the number of independent experiments. One-way ANOVA or two-way ANOVA followed by Dunnetts’ post hoc test were used for the statistical evaluation of experimental data in GraphPad Prism 9.0 (La Jolla, CA, USA). Statistical comparison of excitatory and inhibitory inputs in the spiking models were performed with one-way ANOVA with Tukey’s post hoc test, and for the sensitivity analysis the Kruskal-Wallis test with Dunn’s post hoc test. Sample sizes and P values are shown in figure legends and supplementary Tables A.1 and A.2. Differences between groups were considered statistically significant if P < 0.05 for electrophysiology experiments, and P < 0.001 for modelling results.

## RESULTS

### The clinical phenotype of a patient carrying the recurrent T162I Na_v_1.1 variant

We obtained the clinical data of a female patient carrying a *de novo* T162I variant associated with NDEEMA. The patient was born with bilateral upper limb arthrogryposis (camptodactyly) and congenital left hip dysplasia. She developed tonic seizures associated with apnoea on the second day of life, followed by tonic-clonic seizures on day three. At two months she developed peri-oral myoclonia proceeding to bilateral clonic seizures, and within six months suffered episodes consistent with infantile spasms. There were frequent focal clonic seizures and episodes of focal motor status epilepticus that involved the distal lower limbs, facial musculature, and eyelids. These could last many hours and were triggered by sensory stimuli such as diaper changes. At ∼18 months a hyperkinetic movement disorder emerged which progressed to near-continuous choreo-athetoid movements most prominent in the upper limbs and face. The patient’s development was delayed globally, but by 30 months she reached complete head control, could roll over, grasp and sit with minimal support. Her motor function regressed at 3 years coinciding with an increase in seizure frequency and at 7 years of age she was unable to sit or stand, required head support in a wheelchair, was unable roll over and performed minimal voluntary movements with her arms.

The patient’s seizures were refractory to multiple antiseizure medications (ASMs). Initial treatment with phenobarbital led to a brief period of seizure freedom lasting six weeks. Stiripentol produced a clear increase in seizure frequency which was confirmed with a second exposure. Clobazam, rufinamide, lamotrigine, acetazolamide and chloral hydrate appeared to mildly worsen clonic and myoclonic seizures. Levetiracetam and brivaracetam were well tolerated, but ineffective, whereas sodium valproate, topiramate and lacosamide were both well tolerated and produced reductions in seizure frequency.

### Biophysical properties of Na_v_1.1 variants associated with NDEEMA and EIDEE/MD and their impact on neuronal firing

To characterize the impact of Na_V_1.1 variants upon channel biophysical properties, including the T162I variant described above, we performed voltage clamp experiments on CHO cells transfected with T162I, I236V, P1345S, or R1636Q Nav1.1 channels (Fig. 1). Compared to wild-type, depolarisation-activated sodium current densities of all variants were unchanged (Figs. 1b and 1c, and Table A.1). The activation curves of three variants (T162I, P1345S, R1636Q) were hyperpolarised, suggesting a gain-of-function, or unchanged (I236V). The steady-state inactivation curves were either depolarized (R1636Q) or hyperpolarized (T162I) consistent with gain- or loss-of-function respectively, or unchanged (I236V and P1345S) (Figs. 1d and 1e, and Table A.1). R1636Q channels also recovered from inactivation at a faster rate (Fig. 1h and Table A.1) supporting a gain-of-function effect through increased sodium channel availability. Finally, we also recorded the I_NaP_ associated with each variant, which is an important determinant of the voltage-gated sodium current that contributes to neuronal excitability (Stafstrom, 2007). Compared to wild-type, the I_NaP_ of the I236V, P1345S, and R1636Q variants was increased over a wide range of membrane potentials (Figs. 1f and 1g, and Table A.1), again suggesting a gain-of-function mechanism.

**Figure 1.**
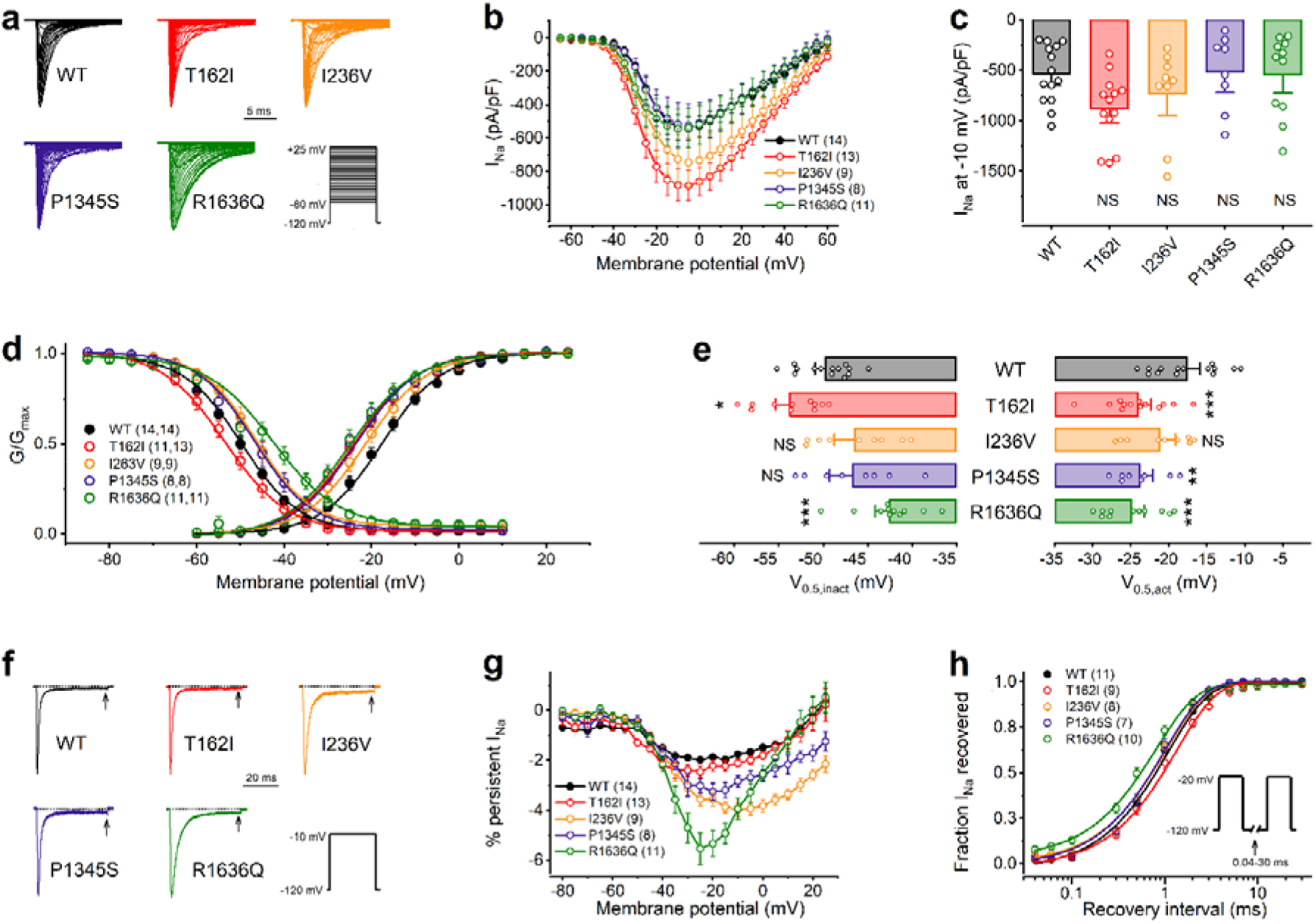
Biophysical properties of bold Na_v_1.1 channel variants associated with early infantile encephalopathy. **a**) Representative families of sodium currents in Chinese hamster ovarian (CHO) cells transiently expressing wild-type (WT), T162I, I236V, P1345S, or R1636Q channels. Currents traces shown were elicited using step depolarizations between −60 and +25 mV in 5 mV increments, from a holding potential of −120 mV (inset protocol). Traces were normalized for the WT inward current peak amplitude recorded at −5 mV. **b**) Current density-voltage relationships. Peak currents were recorded at voltages between −80 to +60 mV in 5 mV increments from a holding potential of −120 mV and were normalized by cell capacitance. WT, black solid circle; mutants, colored open circles. **c**) Mean ± SEM values of the current densities recorded at −10 mV; individual data are shown in front of the bars. **d**) Voltage-dependent activation and inactivation curves, obtained by non-linear least-squares fits of Boltzmann equations to normalized conductance (G/G_max_) data. **e**) Mean ± SEM values of the half-maximal voltages of activation and inactivation (V_0.5,act_ and V_0.5,inact_, respectively) of WT Na_v_1.1 and mutant Na_v_1.1 channels. Individual V_0.5,act_ and V_0.5,inact_ data are shown in front of the bars. Relative to WT, hyperpolarising shifts of the V_0.5,act_ values and depolarising shifts of the V_0.5,inact_ values are consistent with gain-of-function (GoF). The T162I variant exhibits mixed GoF and loss-of-function (LoF) due to the hyperpolarising shifts of the V_0.5,act_ and V_0.5,inact_ values. **f**) Representative normalised Na_v_1.1 current traces of the variants demonstrating the absence or presence of persistent sodium current (I_Na-P_). Current were elicited at −10 mV from a holding potential of −120 mV (inset protocol); dotted lines indicate zero current level; I_Na-P_ was determined 40 ms after the onset of the depolarizing voltage step (arrow). **g**) I_Na-P_-voltage relationships, plotted as percentage of peak sodium current. Currents were elicited using depolarizing voltage steps in 5 mV increments in the voltage range between −80 and +25 mV, from a holding potential of −120 mV. WT, black solid circle; mutants, colored open circles. Data are represented as mean ±SEM. Statistically significant differences between the WT and mutant channels were determined using two-way ANOVA, followed by Dunnett’s post-hoc test. Asterisks indicate ^*^P < 0.05. The statistical evaluation of the I_Na-P_ values elicited at −20 mV are shown in Table A.1. **h**) Recovery from fast inactivation was assessed using a double-pulse voltage protocol (inset). The fraction of current recovered during the second test pulse was plotted against a range of recovery interval (0.04-30 ms). The time constant (τ) of recovery was estimated by fitting a single exponential function to the data (see Methods). WT, black open bar; mutant, colored open bars. Data in b, d, g, and h represent mean ± SEM values; n values for all variants are shown in parentheses. See the statistical evaluation of biophysical parameters in Table A.1. ^*^P < 0.05, ^**^P < 0.01, and ^***^P < 0.001, one-way ANOVA, followed by Dunnett’s post-hoc test. NS, statistically not significant difference compared to WT.

The relationship between changes in Na_V_1.1 biophysical properties and neuronal excitability can be counterintuitive (Berecki et al., 2019). Therefore, we used the DAPC approach (Berecki et al., 2019; Berecki et al., 2018) to predict the impact of each *SCN1A* variant on the firing activity of a single-compartment neuron model (Fig. 2). Here, the heterologously expressed Na_v_1.1 current was implemented as an external input conductance resulting in a hybrid experimental-computational neuron model. Hybrid neurons incorporating the I236V, P1345S, or R1636Q conductance exhibited significantly higher peak firing rates relative to wild-type, and the presence of the T162I and R1636Q conductance produced a lower firing threshold (rheobase) (Fig. 2c and Table A.2). Indeed, incorporating the R1636Q conductance produced spontaneous firing at baseline without external current input. Interestingly, the T162I conductance led to depolarization block at higher input amplitudes, similar to the previously described T226M *SCN1A* variant (Berecki et al., 2019) (Figs. 2a and 2b). Depolarization block in the presence of T162I variant is consistent with the hyperpolarizing shift in voltage dependence of inactivation, and results in action potential collapse at stimulus levels where normal firing is seen with WT. Our data also suggest that the presence of increased I_NaP_ measured over a wide voltage range for I236V, P1345S, and R1636Q relative to WT (Fig. 1g), is a key determinant of a sustained hybrid neuron firing phenotype.

**Figure 2.**
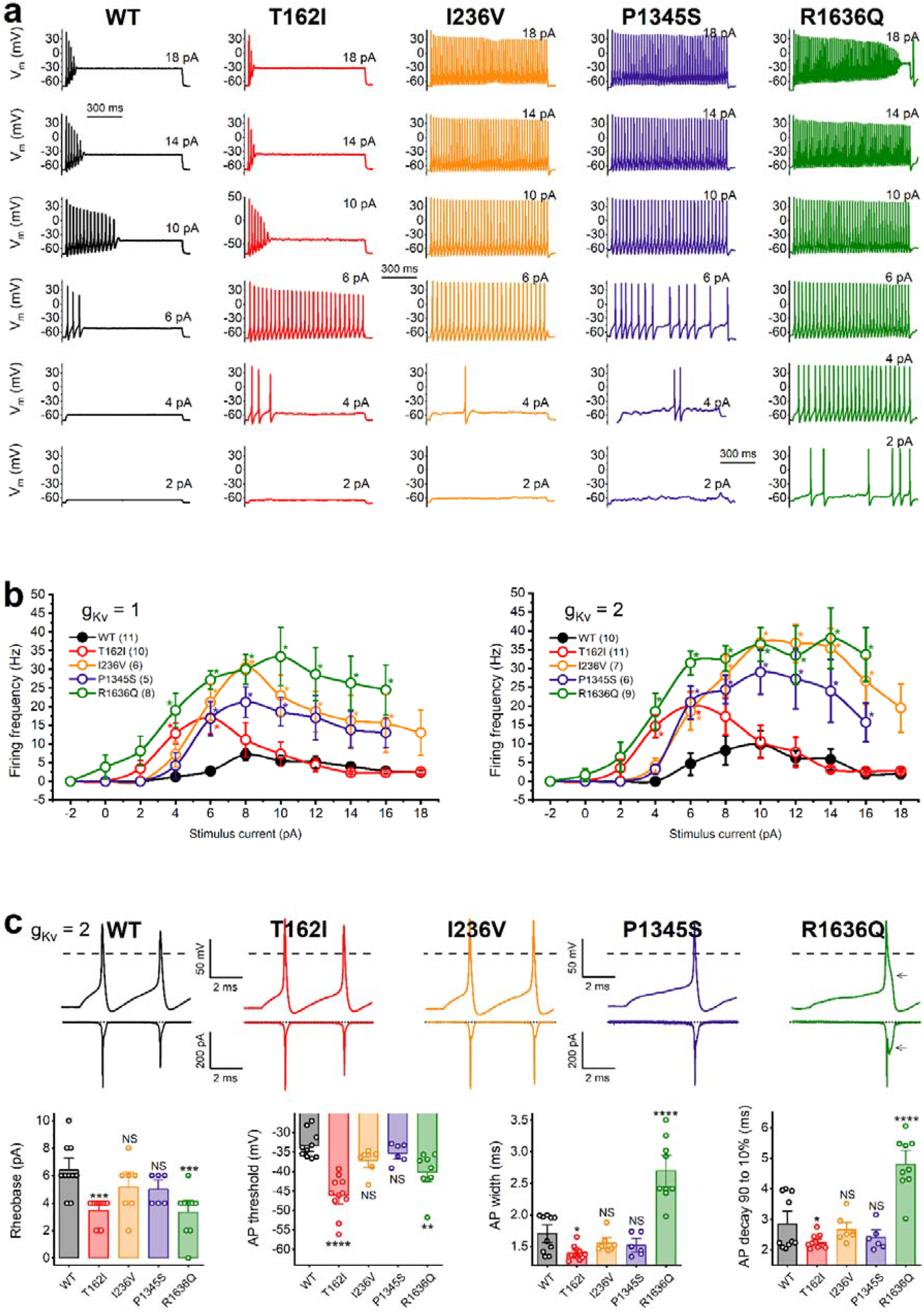
The impact of the wild-type (WT), T162I, I236V, P1345S, or R1636Q Na_v_1.1current on the excitability of axon initial segment (AIS) compartment model in dynamic action potential clamp experiments. **a**) Representative action potential firing with WT, I236V, P1345S, or R1636Q variant in response to 2, 4, 6 10, 14, and 18 pA step current stimuli. The in silico Na_v_1.6 conductance (gNa_v_1.6) was set to zero, whereas the in silico potassium channel conductance (gKv) was adjusted to 200% of the original gKv (gNa_v_1.6 = 0, gK_v_ = 2, respectively). **b**) Input-output relationships with gK_v_ = 1 (left) or gK_v_ = 2 (right) settings. Action potential firing was elicited by current steps in 2 pA increments in the range between −2 and 18 pA. Data are represented as mean ±SEM; n values, the number of independent experiments, are shown in parentheses. WT, black solid circle; mutants, colored open circles. Statistically significant differences between the wild-type (WT) and mutant channels were determined using two-way ANOVA, followed by Dunnett’s post-hoc test; asterisks indicate ^*^P < 0.05. The individual data points and the results of statistical evaluation are included in Table A.2. **c**) Representative action potentials (gK_v_ = 2) (upward deflections) and the external input Na_v_1.1 variant currents (downward deflections), contributing current to the action potentials in the AIS hybrid neuron. Dashed lines indicate 0 mV, dotted lines indicate 0 pA. The first action potential elicited by a current step 2 pA above the rheobase was analysed to determine selected action potential characteristics, including action potential threshold, action potential width and action potential decay. Bars show the mean ± SEM; individual data are shown in front of the bars; n values are shown in b. Differences in action potential characteristics were assessed using one-way ANOVA followed by Dunnett’s post-hoc test (^*^P < 0.05, ^**^P < 0.01, ^***^P < 0.001, and ^****^P < 0.0001); NS, statistically not significant difference compared to WT. The statistical evaluation of the action potential characteristics is shown in Table A.2.

To further highlight the functional consequences of the variants, we also analysed action potential morphology in DAPC experiments. Action potential width decreased in the presence of T162I, whereas R1636Q resulted in both increased action potential width and decay time compared with wild-type (Fig. 2c and Table A.2). In summary, DAPC experiments demonstrate that all four *SCN1A* variants associated with NDEEMA and EIDEE/MD increase the excitability of hybrid neuron models through decreased neuronal rheobase and/or higher peak firing rates.

### Enhanced interneuron excitability can promote network instability through homeostatic plasticity

Due to the preferential expression of *SCN1A* within inhibitory interneurons, our results suggest that *SCN1A* gain-of-function variants associated with NDEEMA and EIDEE/MD would be expected to enhance cortical inhibition and suppress hyper-excitable states such as epilepsy. However, brain networks can undergo homeostatic plasticity to restore normal levels of activity that support cortical processing (Hengen et al., 2013; Keck et al., 2017), and it is possible that such adaptations may produce unexpected consequences upon network function. Several different mechanisms can contribute to homeostatic plasticity to restore pyramidal cell (PC) firing rates in-vivo, including scaling of excitatory synaptic strength (Keck et al., 2017; Turrigiano et al., 1998), scaling of inhibitory synaptic strength (D’Amour J and Froemke, 2015; Hennequin et al., 2017; Vogels et al., 2011; Xue et al., 2014) and upregulation of intrinsic neuron excitability (Gasselin et al., 2015; Maffei and Turrigiano, 2008). Indeed, these mechanisms have been observed in epilepsy models suggesting they may play an important role in epileptogenesis (Lignani et al., 2020; Nasrallah et al., 2022; Zhang et al., 2021).

Therefore, to explore the impact of *SCN1A* gain-of-function and homeostatic plasticity within a neuronal network, we used a spiking cortical network model containing an excitatory (PC) and inhibitory (PV) population (Bryson et al., 2021). *SCN1A* gain-of-function was first introduced by enhancing PV interneuron excitability through a reduction of firing threshold (Figs. 3a and 3b) which suppressed PC firing rates (Supplementary Fig. A.1). We then developed three models that incorporated different forms of homeostatic plasticity to restore baseline PC firing rates (Figs. 3c and 3d): a reduction of PV-to-PC synaptic strength (i.e., inhibitory synaptic plasticity, Model 1), an increase in PC-to-PC synaptic strength (i.e., excitatory synaptic plasticity, Model 2) and an increase of intrinsic PC excitability (Model 3).

**Figure 3.**
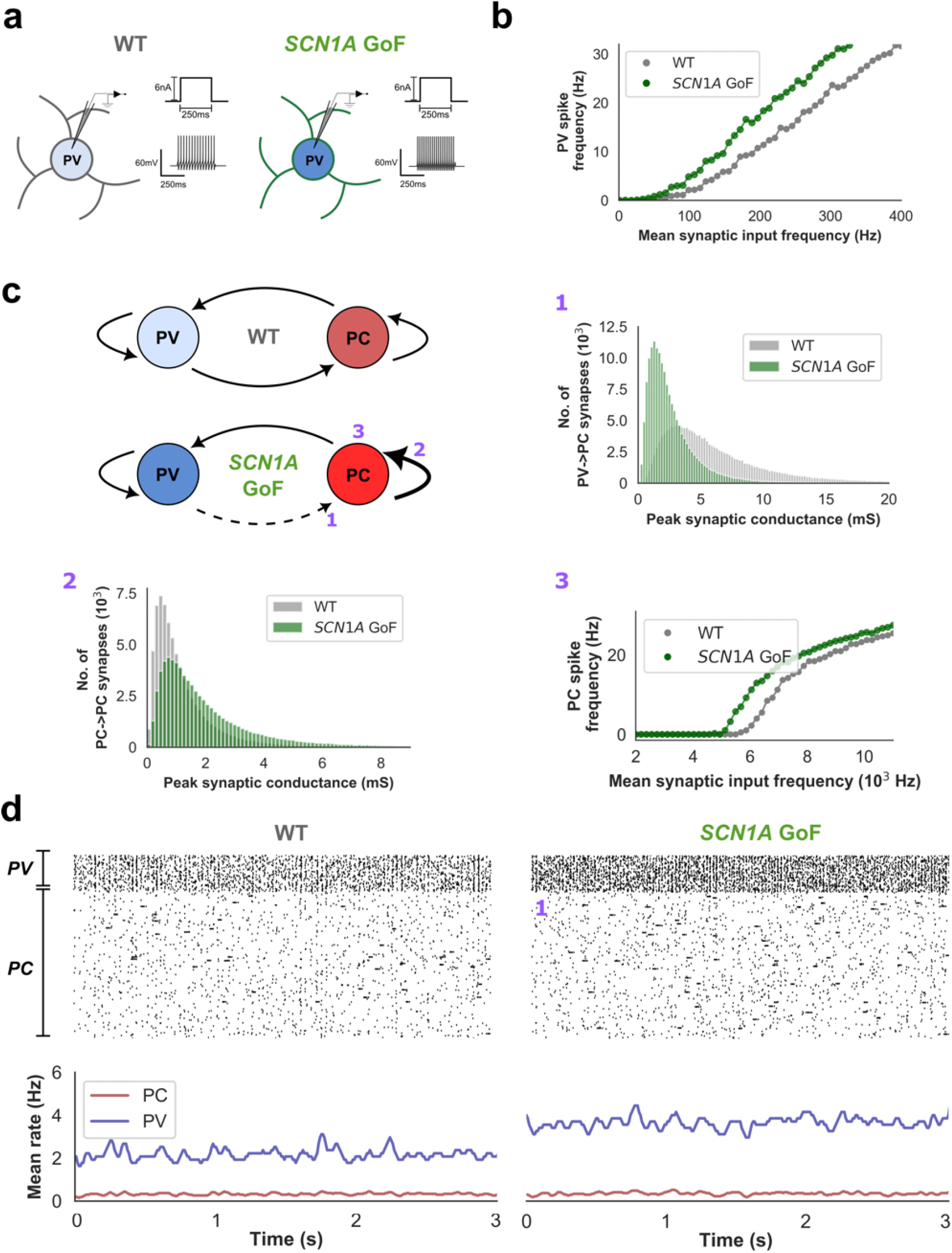
Baseline characteristics of the wild-type and *SCN1A* gain-of-function network models. *SCN1A* gain-of-function (GoF) was modelled by increasing the intrinsic excitability of PV inhibitory interneuron models, resulting in higher spike frequencies in response to constant current input (**a**) or synaptic inputs (**b**). **c**) Schematic representation of the network models containing excitatory (PC) and either WT (upper) or *SCN1A* GoF (lower) PV interneuron populations. To model homeostatic plasticity in the presence of *SCN1A* GoF, PC firing rates were restored to WT levels by using Model 1, Model 2, or Model 3 (see details in the text). **d**) Raster plots (upper) and mean firing rates (lower) of the PC and PV population in the WT and *SCN1A* GoF networks (Model 1) during baseline spiking. At baseline, the PV population has a higher mean firing rate in the *SCN1A* GOF network due to increased intrinsic excitability (3.7 ± 1.2 Hz vs 2.2 ± 1.1 Hz over the 3s simulation shown, *P* < 0.001, Welch’s t-test). In contrast, PC firing rates are similar in both models (0.34 ± 0.2 Hz and 0.36 ± 0.17 Hz over the 3s simulation shown, N.S., Welch’s t-test) due to the compensatory reduction of PV-to-PC synaptic strength. Models 2 and 3 possess similar baseline firing characteristics (Supplementary Figure A.1).

The stability of the wild-type and *SCN1A* gain-of-function networks was then explored by applying a brief external stimulus to the PC population (Fig 4a). Interestingly, despite similar baseline PC firing rates in all networks, we observed variability in response to external perturbation (Figs. 4b and 4c). Although Models 2 & 3 exhibited a similar network response to wild-type, Model 1 generated bursts of synchronous activity suggesting an important role for PV-to-PC synaptic strength for maintaining network stability. We also assessed the relative contribution of intrinsic PC excitability and PC-to-PC synaptic strength to network instability by increasing each parameter until synchronous activity emerged following external stimulation (Supplementary Fig. A.2). Here, we found that large increases of PC-to-PC synaptic strength could predispose to network instability, but that enhancing intrinsic PC excitability to an extent that produced similar firing rates did not. By measuring excitatory and inhibitory currents onto PC neurons in each Model, we found that only Models 1 and 2 led to an increase in EI ratio, driven by a reduction and increase in feedback inhibition, respectively, and an increase in recurrent excitation (Model 2) (Figs. 4d-4f).

**Figure 4.**
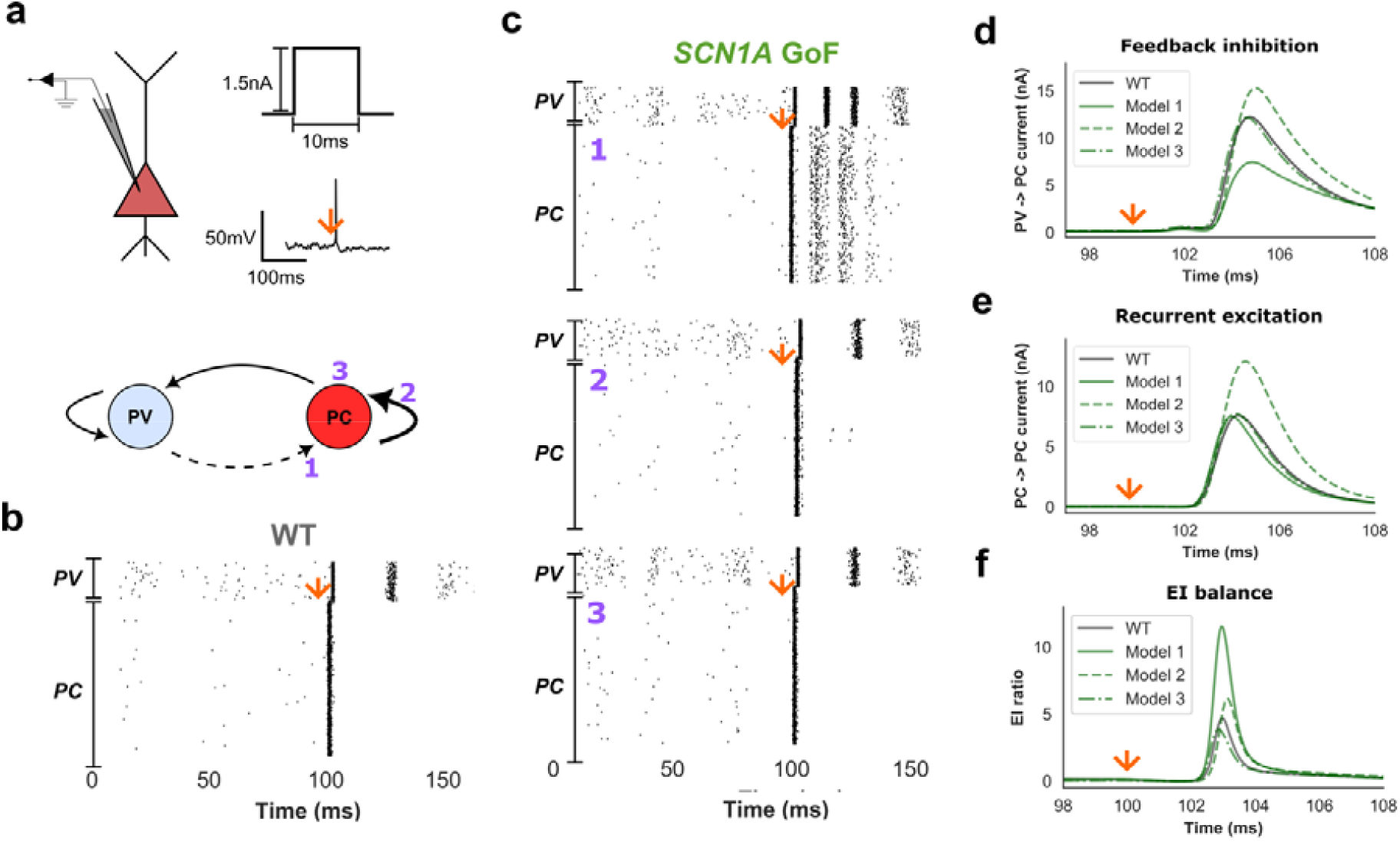
Response of the wild-type and *SCN1A* GoF networks to external stimulation. **a**) To explore network stability, neuron models within the PC population were stimulated with a brief current that elicited a single spike. This stimulus was applied to WT and all *SCN1A* GoF models. **b, c**) Raster plots of WT and *SCN1A* GoF networks following stimulation of the PC population (orange arrow) at 100 ms. Compared to the WT network and Models 2 and 3, Model 1 was more sensitive too external perturbation and exhibited recurrent bursts of population activity. Similar findings were observed in Model 2 if greater increases in PC-to-PC synaptic weights were introduced (Supplementary Fig. A.2). In contrast, enhancing intrinsic PC excitability alone did not predispose to recurrent activity (Supplementary Fig. A.2). Feedback inhibition onto the PC population (**d**), recurrent PC-to-PC excitation (**e**) and network EI balance (**f**) following external stimulation. Reduced PV-to-PC synaptic strength (Model 1) led to a reduction of peak feedback inhibition compared to WT (7.5 ± 0.1 nA vs 12.2 ± 0.12 nA, *P* < 0.001, one-way ANOVA with post-hoc Tukey’s test), whereas increased PC-to-PC synaptic strength (Model 2) produced a large increase of recurrent excitation (12.1 ± 0.2 nA vs 7.6 ± 0.1 nA, *P* < 0.001). Both Model 1 and 2 had increased peak EI ratio compared to WT (10.5 ± 0.3 and 4.8 ± 0.2, respectively, vs 3.5 ± 0.1, *P* < 0.001).

It is possible that different homeostatic plasticity mechanisms may interact to restore network firing rates. Therefore, to explore the impact of these interactions and to further elaborate the importance of each mechanism upon network stability, we performed a sensitivity analysis by incrementally varying each parameter and then re-assessing the network response to external stimulation (Figs. 5 a and b). We found that reduced PV-to-PC synaptic strength and increased PC-to-PC synaptic strength predisposed to a greater network response (Fig 5c). In contrast, increasing PC excitability did not significantly enhance network response. Overall, these findings suggest that while several homeostatic plasticity mechanisms may act to restore population firing rates, the relative contribution of each mechanism can drive distinct network behaviour that differentially predisposes to network instability, and which may also account for phenotypic variability in the presence of *SCN1A* variants (Fig 5d).

**Figure 5.**
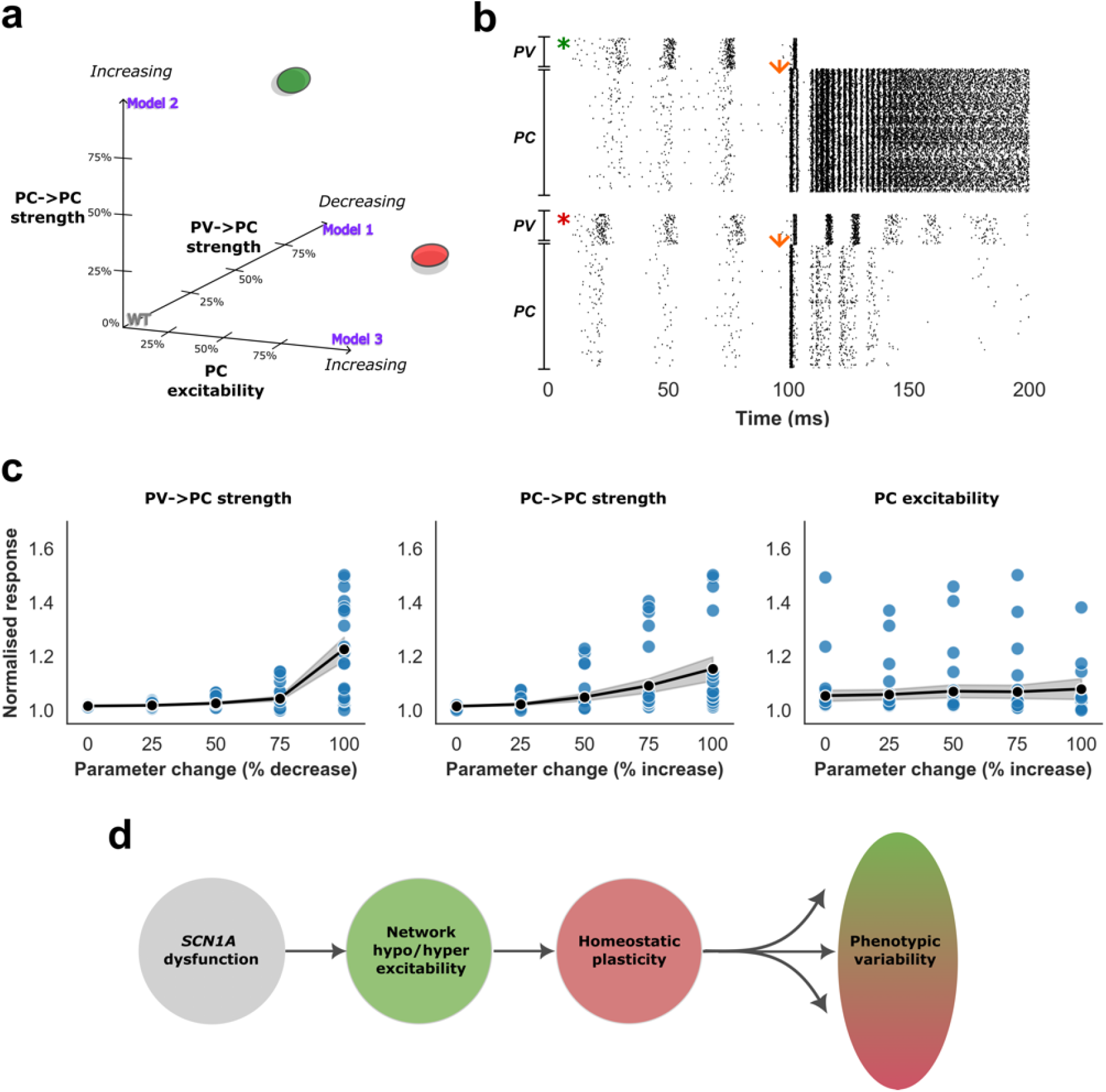
Contribution of homeostatic plasticity mechanisms to network stability. **a**) To investigate the relative contribution and interactions between homeostatic plasticity mechanisms to network instability, a sensitivity analysis was performed by modulating each parameter associated with Models 1, 2, and 3 by 25% (i.e., 125 models in total). Here, 100% denotes the parameter value from Figs. 3 and 4 that restore baseline PC firing rates. Reducing PV-to-PC and increasing PC-to-PC synaptic strength to 100% (green circle) generated recurrent sustained activity in response to external stimuli, demonstrated in panel b. In contrast, reducing PV-to-PC and enhancing intrinsic PC excitability generated brief synchronous activity similar to Fig. 4c, suggesting that intrinsic PC excitability exerts less influence upon network stability. **c**) Parameter change vs total spikes generated within 10 ms of stimulus (normalised to the WT model) for all 125 models. Reduced PV-to-PC and increased PC-to-PC synaptic strength exerted a significant increase in network response (1.02 ± 0.003 vs 1.23 ± 0.23 and 1.02 ± 0.005 vs 1.15 ± 0.22, respectively for 0 % vs 100 % parameter change, P < 0.001, Kruskal-Wallis Test with post-hoc Dunn test) whereas changes to intrinsic PC excitability did not significantly modify the network response (1.05 ± 0.1 vs 1.08 ± 0.19 for 0 % vs 100 % parameter change, P = NS). **d**) We propose that an important contributor to phenotypic variability relates to different homeostatic plasticity mechanisms that interact to restore basal population firing rates.

Finally, we investigated why changes to synaptic strength (Models 1 & 2) may predispose to network instability using a simplified model of mean firing rates with an excitatory (E) and inhibitory (I) population, and under the assumption of linear input-output relationships (Figs. 6a-c). In this model, due to linearity of the population response, changes to threshold shift the steady-state firing rates but do not scale the strength of feedback inhibition or recurrent excitation, and stability is preserved (Supplementary Figs. A.2c and 2d). In contrast, these properties are scaled with reduced I-to-E and increased E-to-E synaptic strength, predisposing to network instability (Fig. 6d) and sensitivity to external stimuli (Fig. 6e), similar to the spiking network model.

**Figure 6.**
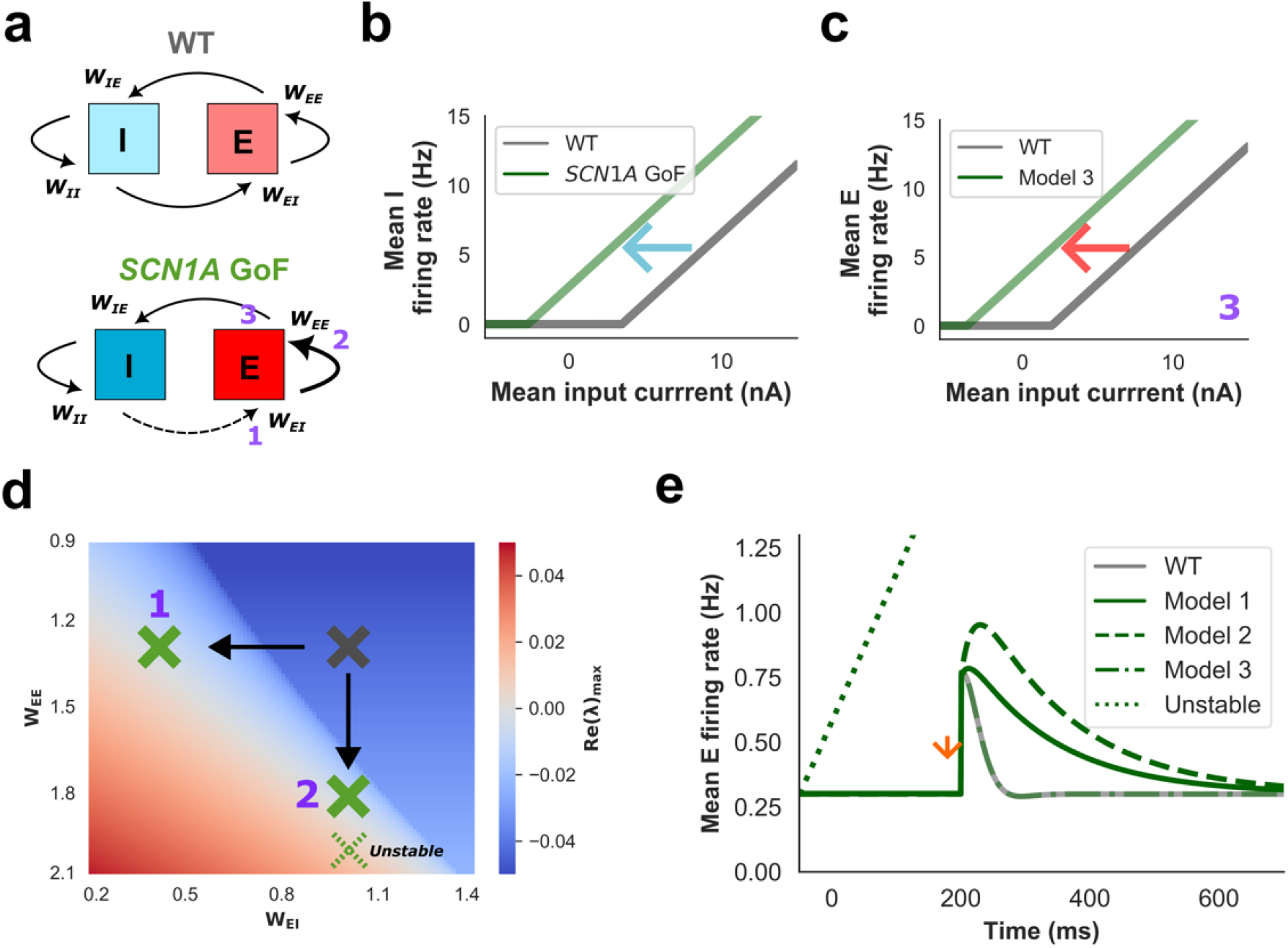
Impact of homeostatic plasticity in a simplified firing-rate model. The model contains an excitatory (E) and inhibitory (I) population (**a**) with linear input-mean firing rate relationships. *W*_*EI*_ and *W*_*IE*_ denote inhibitory-to-excitatory and excitatory-to-inhibitory synaptic strength, respectively; *W*_*EE*_ and *W*_*II*_ denote excitatory-to-excitatory and inhibitory-to-inhibitory synaptic strength, respectively. *SCN1A* GoF was modelled by first reducing the firing threshold of the inhibitory population (**b**) and then either reducing *W*_*EI*_ synaptic strength (Model 1), increasing *W*_*EE*_ synaptic strength (Model 2), or reducing the threshold of E firing rate (Model 3, **c**) to restore the baseline E firing rates. **d**) Changes to either *W*_*EE*_ or *W*_*EI*_ modify network stability, demonstrated by the real value of the eigenvalue of highest value associated with the model’s Jacobian matrix (purple numbers denote model and red regions unstable parameter values**). e**) Changes to model stability are also reflected by the response to external stimulation (orange arrow) of the E population. Models 1 and 2 exhibit a sustained response whereas Model 3 is unchanged.

## DISCUSSION

*SCN1A* gain-of-function variants have been associated with a spectrum of neurological disorders, including early onset DEE, arthrogryposis multiplex congenita and/or movement disorder (Berecki et al., 2019; Brunklaus et al., 2022), but the pathophysiological mechanisms that give rise to the clinical phenotype are poorly understood. In a previous study, we found that model inhibitory interneurons incorporating the T226M variant exhibited depolarisation block with strong current input, suggesting a cellular mechanism through which *SCN1A* gain-of-function may predispose to cortical disinhibition underlying early onset DEE (Berecki et al., 2019). However, it is unclear if this mechanism applies to other *SCN1A* gain-of-function variants. Furthermore, the potential impact of these changes upon neuronal network activity has not been characterized.

In this study, we report a recurrent T162I variant identified in a patient with NDEEMA, further corroborating genotype-phenotype associations. We then characterized the biophysical properties of four *SCN1A* variants associated with NDEEMA or EIDEE/MD and assessed their impact upon the excitability of hybrid neuron models reproducing the properties of PV interneurons. All variants predisposed to reduced neuronal rheobase and/or higher peak firing rates compared to wild type, consistent with a gain-of-function effect at the neuronal scale. Finally, we incorporated PV interneuron hyperexcitability into a cortical network model and then introduced three simple forms of homeostatic plasticity that act to restore PC firing rates. We found that homeostatic plasticity mechanisms exert differential effects upon network stability, with changes to synaptic weight predisposing to synchronous excitatory activity.

Previous work by ourselves and other groups (Berecki et al., 2019; Brunklaus et al., 2022; Clatot et al., 2022; Matricardi et al., 2023) suggest that early onset DEE *SCN1A* variants show unaltered current densities in transfected mammalian cells compared to wild-type. Therefore, the gain-of-function effect of these variants upon neuronal function may instead be attributable to other biophysical characteristics (Fig. 1). For example, we observed a hyperpolarizing shift in the voltage dependence of activation across three variants (T162I, P1345S and R1636Q) suggesting a lower threshold for action potential generation. Likewise, other groups also showed a hyperpolarising shifts in the voltage dependence of activation for T162I (Matricardi et al., 2023) and R1636Q (Clatot et al., 2022). Our results also confirm the I_NaP_ increase for R1636 and I236V, as shown in previous studies (Brunklaus et al., 2022; Clatot et al., 2022). Nevertheless, some of the biophysical characteristics of Nav1.1 variants assessed in this study and previously also show considerable variability (Supplementary Table A.3). A gain-of-function effect is supported by our DAPC experiments that demonstrate a lower firing threshold compared to wild-type for the T162I and R1636Q variants (Fig. 2). Furthermore, our use of DAPC also revealed additional mutation-specific biophysical consequences. Hybrid neurons incorporating the T162I variant exhibited depolarisation block and loss of action potential firing with high intensity stimuli, an effect reminiscent of the mixed biophysical defects in T266M Na_v_1.1 channels associated with early infantile DEE (Berecki et al., 2019), whereas I236V, P1345S, and R1636Q variants had higher and sustained firing rates relative to wild-type in response to high intensity stimuli. This variable impact upon neuronal excitability may relate to difference in channel gating biophysics. The presence of hyperpolarising and depolarising shifts of steady-state activation and inactivation, respectively, and enhanced I_NaP_ together enhance the ‘window’ current generated by voltage-gated sodium channels, and would be expected to increase peak firing rates (I236V, P1345S and R1636Q). In contrast, hyperpolarising shifts of both activation and inactivation (T162I) may predispose to depolarisation block, as previously demonstrated (Berecki et al., 2019).

Epilepsy is considered a disorder of abnormal cortical excitation. In Dravet syndrome associated with *SCN1A* loss-of-function, PV interneuron dysfunction is thought to predispose to disinhibition and cortical pyramidal cell hyperexcitability leading to an epileptic phenotype (Catterall et al., 2010; Zuberi et al., 2011). Furthermore, other interneuron subtypes such as SST interneurons are also implicated in the pathogenesis of Dravet Syndrome and may exert a synergistic effect (Tai et al., 2014). As such, it is counterintuitive that *SCN1A* gain-of-function is also associated with epilepsy. A high EI ratio may predispose to seizures, whereas excess inhibition may suppress PC firing rates and impair normal cortical function and maturation, and it has been shown that homeostatic plasticity allows cortical networks to restore PC firing rates and EI balance (Keck et al., 2017; Lignani et al., 2020; Sutton and Schuman, 2006; Turrigiano, 2008). Although homeostatic plasticity is thought to be important in the pathogenesis of epilepsy, but it has not yet been studied in detail (Lignani et al., 2020). Nevertheless, several studies have revealed homeostatic regulation of voltage-dependent conductance’s that modulate intrinsic neuron excitability. For example, activity-dependent modulation of sodium (Aptowicz et al., 2004; Desai et al., 1999), potassium (Cudmore et al., 2010), calcium (Desai et al., 1999), and HCN channels (Gasselin et al., 2015) has been demonstrated, and scaling of Kv1.1 subunit transcription has been associated with homeostatic plasticity in temporal lobe epilepsy (Kirchheim et al., 2013). Furthermore, complex changes to synaptic strength have been observed in *SCN1A*^+/-^ mice that may help to restore PC firing rates (De Stasi et al., 2016), and strengthening of recurrent excitatory synapses can contribute to epileptogenesis in temporal lobe epilepsy (Nasrallah et al., 2022).

Here, using a spiking network model, we show that homeostatic plasticity mechanisms that restore PC firing rates can exert differential impact upon network stability, and may predispose to epileptiform-like activity. Our findings raise the possibility that an important driver of phenotypic variability and treatment response in *SCN1A* disorders could relate to the compensatory homeostatic mechanisms that are recruited to recover normal cortical firing rates. Furthermore, our findings suggest that changes to synaptic strength may be of particular importance for the emergence of cortical hyper-excitability. These insights may carry therapeutic implications as selective modulation of homeostatic pathways that drive the network into a less excitable state may be of particular benefit for the treatment of epilepsy (Lignani et al., 2020).

Although our model proposes a mechanism for how changes to intrinsic neuron function may produce unexpected effects upon network activity, there are limitations to this approach and our predictions require experimental validation in an *SCN1A* gain-of-function mouse model. To simplify our analysis and provide a proof-of-principle we have used a network model with a single homogenous interneuron population with firing characteristics typical for PV-positive interneurons, and additional complexity is anticipated if other interneuron subtypes are also included. Furthermore, we modelled both *SCN1A* gain-of-function and a homeostatic increase in PC excitability (Model 3) by reducing neuronal threshold, and it is possible that *SCN1A* variant-induced and homeostatic adaptations to intrinsic voltage-dependent conductances may modify other neuronal properties such as gain. Gain modulation can exert different effects upon network behaviour and can strengthen recurrent excitatory activity to enhance population response (Bryson et al., 2020; Ferguson and Cardin, 2020; Haider and McCormick, 2009; Rubin et al., 2015). Finally, our model may imply that *SCN1A* loss-of-function associated with Dravet Syndrome could produce compensatory changes to synaptic strength that would be expected to reduce network stability. However, it remains unknown which homeostatic mechanisms are recruited under different conditions, and it is likely there exists a range over which synaptic strength can adjust for changes to intrinsic neuron excitability. Therefore, if either channel gain- or loss-of-function exceeds this compensatory ‘envelope’ a hyper-excitable phenotype may emerge.

*SCN1A* gain-of-function variants are also associated with familial hemiplegic migraine type 3 (FHM3) (Auffenberg et al., 2021). Similar to variants associated with DEE, FHM3 *SCN1A* gain-of-function variants arise through several mechanisms, including increased I_NaP_ and delayed recovery from inactivation (Kahlig et al., 2008; Mantegazza and Cestele, 2018). Missense mutations associated with FHM3 can result in folding defects and reduced channel expression, yet even partial expression of such variants may be sufficient for overall gain-of-function biophysical characteristics (Cestele et al., 2013; Dhifallah et al., 2018). Interestingly, epilepsy and migraine share certain pathophysiological similarities: for example, depolarisation block due to sodium channel inactivation in PCs has been observed in both conditions, resulting in a hyperexcitable state that propagates through cortical networks. However, the speed and dynamics of propagation differ significantly between these two conditions, and it remains unknown how gain-of-function in the same gene can result in such distinct clinical phenotypes (Mantegazza and Cestele, 2018). In a cortical spreading depression (CSD) model for FHM3 due to a Nav1.1 gain-of-function mutation, the early phase of CSD was associated with the increased activity of fast-spiking interneurons and the subsequent increase of extracellular K^+^ concentration in brain slices (Auffenberg et al., 2021). Similarly, CSD could be initiated in the absence of GABAergic or glutamatergic synaptic transmission by an increase in the activity of interneurons resulting in extracellular K^+^ build (Chever et al., 2021). In an FHM3 computational microcircuit model, Nav1.1 channel gain-of-function in interneurons led to increased K channel activation, CSD, and modification of ion fluxes (Lemaire et al., 2021). K^+^ accumulation in these models suggest that besides the increased activity of interneurons other cellular mechanisms are also involved in FHM3. Our network does not specifically changes in ionic fluxes, but involvement of such mechanisms in early onset SCN1A DEE cannot be excluded. These questions may be addressed through generating animal models of *SCN1A* gain-of-function variants. Although it has been proposed that FHM3 variants may cause larger functional effects than early onset DEE variants (Brunklaus et al., 2022) this hypothesis requires the analysis of a larger set of *SCN1A* gain-of-function variants associated with DEE or FHM3.

In conclusion, our findings support previous observations that *SCN1A* gain-of-function can lead to DEE, and we extend these other results by demonstrating enhanced excitability within hybrid models of PV interneurons using DAPC. Furthermore, we propose a mechanism through which homeostatic mechanisms that restore firing rates may predispose to phenotypic variability and a hyper-excitable network state. A crucial goal of future studies is to test these predictions relating to early onset DEE-associated *SCN1A* gain-of-function variants using patient-derived stem cell neurons and animal models. A deeper understanding of homeostatic pathways may reveal novel opportunities for novel and more effective therapeutic interventions.

## Glossary

ASM: anti-seizure medication;
DEE: developmental and epileptic encephalopathy;
DAPC: dynamic action potential clamp;
LoF: loss-of-function;
GoF: gain-of-function

## Acknowledgements

We are grateful to all patients and their families participating in this research. This study was supported by an Australian Research Council Centre of Excellence for Integrative Brain Function grant (CE14010007), National Health and Medical Research Council (NHMRC) program grant (10915693) to S.P., Medical Research Future Fund Genomic Health Futures Mission Project Grant to G.B. and S.P. S.P. is supported by an NHMRC Fellowship. G.B. was partly funded by RogCon, Inc.

## Author contributions

S.P., G.B, and A.B. designed the study, G.B. carried out the electrophysiological studies and analysed and interpreted data. A.B. carried out the in-silico modelling and analysed and interpreted data. T.P. collected and analysed clinical data. G.B., A.B., T.P, and S.P. wrote the manuscript. All authors contributed to reviewing and revising the manuscript.

## Data availability

All data will be available from the corresponding authors upon reasonable request.

