## Supplemental information for "Biophysical characterization and modeling of *SCN1A* gain-of-function predicts interneuron hyperexcitability and a predisposition to network instability through homeostatic plasticity"

___________________________________________________________________________

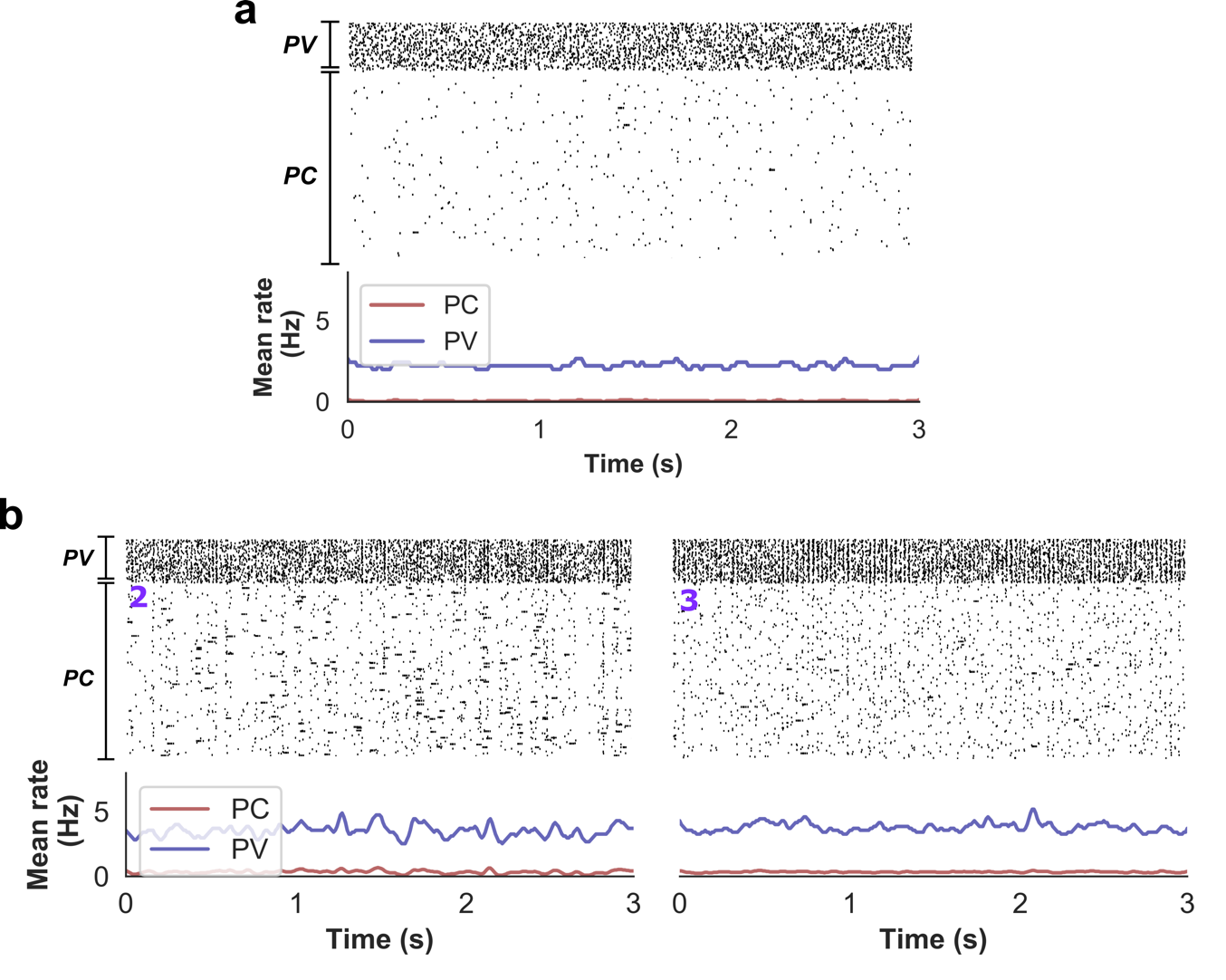

**Figure A.1.** Modulation of the *SCN1A* gain-of-function cortical network model activity by homeostatic plasticity. **a**) Raster plot of network activity after increasing intrinsic parvalbumin positive (PV) interneuron excitability but without introducing homeostatic plasticity. Pyramidal cell (PC) firing rates are suppressed compared to WT (0.04 ± 0.03) and PV firing rates increased (2.25 ± 0.16). Rates expressed as mean ± std. **b**) Raster plots after incorporating increased PC-to-PC synaptic strength (Model 2) and increased intrinsic PC excitability (Model 3) to restore PC firing rates. See raster plot after implementing reduced PV-to-PC synaptic strength (Model 1) in the main manuscript (Fig.3).

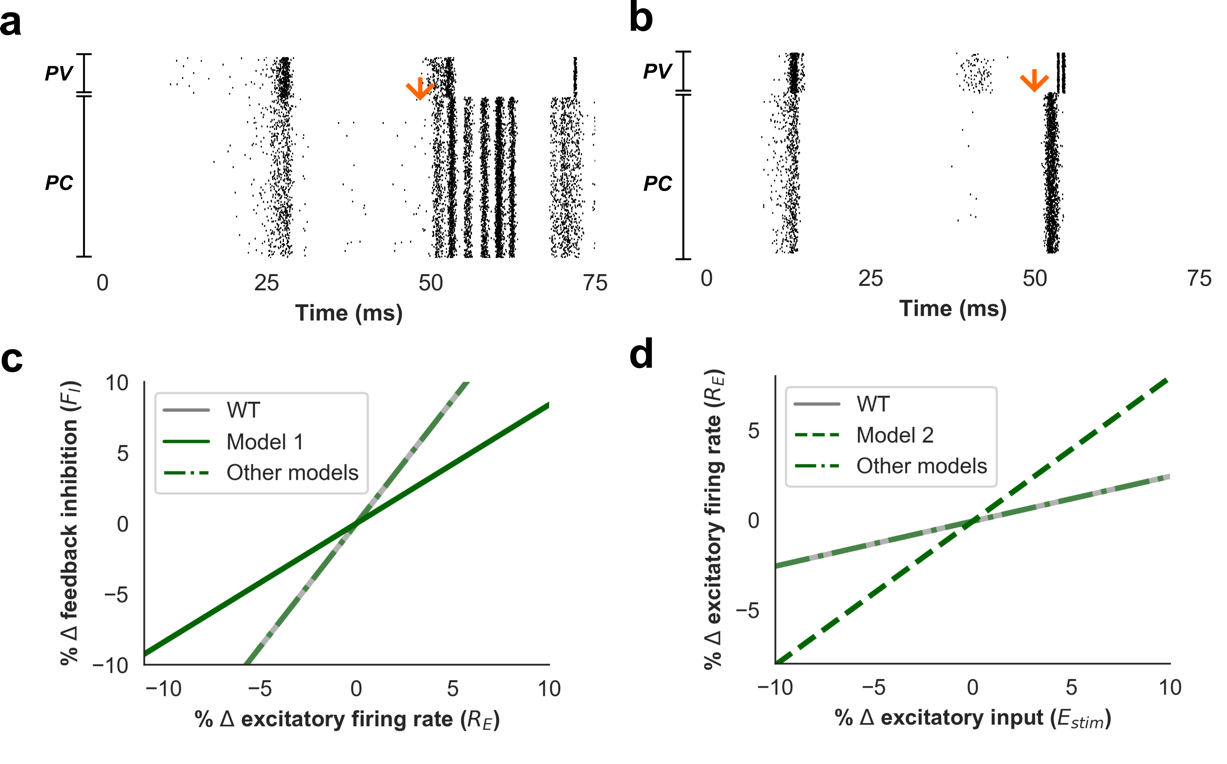

**Figure A.2.** Raster plots of network activity in response to external stimulation at 50 ms after increasing PC-to-PC synaptic weights (**a**) and intrinsic PC excitability (**b**) to values that generate baseline PC firing rates of ~4Hz. Enhancing PC-to-PC synaptic strength, but not intrinsic PC excitability, predisposes to synchronous network activity. **c**) Change in magnitude of feedback inhibition in response to an increase in excitatory firing rate in the simple firing rate models. Model 1 is associated with a deficit of feedback inhibition. **d**) Change in excitatory firing rate in response to external stimuli in the simplified firing rates models, under the assumption of a fixed inhibitory firing rate. Model 2 is associated with greater recurrent excitation leading to a higher change in firing rate.

| Variant | Biophysical property | | | | |
| --- | --- | --- | --- | --- | --- |
|  | **I_Na_ density**  **at −10 mV (pA/pF)** | **V_0.5,act_**  **(mV)** | **V_0.5,inact_**  **(mV)** | **I_NaP_ at −20 mV (% of peak)** | **τ recovery**  **(ms)** |
| WT  n  P value | 535.22 ± 76.9  14  - | −17.63 ± 1.18  14  - | −49.74 ± 0.79  14  - | 1.86 ± 0.15  14  - | 1.08 ± 0.13  11  - |
| T162I  n  P value | 880.38± 95.62^NS^  13  0.059 | −24.02 ± 1.11^***^  13  0.0005 | −53.74 ± 1.07^*^  11  0.036 | 2.22 ± 0.22^NS^  13  0.787 | 1.34 ± 0.08^NS^  9  0.135 |
| I236V  n  P value | 726.59 ± 147.13^NS^  9  0.573 | −21.20 ± 1.06^NS^  9  0.117 | −46.49 ± 1.51^NS^  9  0.179 | 3.56 ± 0.28^***^  9  0.0011 | 1.04 ± 0.08^NS^  8  0.994 |
| P1345S  n  P value | 518.03 ± 132.14^NS^  8  0.999 | −23.80 ± 1.14^*^^*^  8  0.0039 | −46.67 ± 1.75^NS^  8  0.249 | 3.25 ± 0.33^*^  8  0.012 | 1.01 ± 0.07^NS^  7  0.961 |
| R1636Q  n  P value | 544.11 ± 119.28^NS^  11  >0.999 | −24.95 ± 1.15^***^  11  0.0001 | −42.55 ± 1.08^***^  11  0.0001 | 5.32 ± 0.51^****^  11  <0.0001 | 0.81 ± 0.04^NS^  10  0.1001 |

**Table A.1.** Biophysical characteristics of the Na_v_1.1 channel variants in voltage clamp experiments.

Data are represented as mean ± SEM. Statistically significant differences between the wild-type (WT) and mutant channels were determined using one-way ANOVA, followed by Dunnett’s post-hoc test (^*^P < 0.05, ^**^P < 0.01, ^***^P < 0.001, and ^****^P < 0.0001); NS, statistically not significant difference compared to WT. Abbreviations: I_Na_, sodium current; I_NaP_, persistent sodium current; n, number of independent experiments; τ recovery, time constant of recovery from fast inactivation; V_0.5,act_, membrane potential for half-maximal activation;V_0.5,inact_, membrane potential for half-maximal inactivation.

**Table A.2.** Action potential characteristics of the axon initial segment neuronal model incorporating wild-type or mutant Na_v_1.1 channels in dynamic action potential clamp experiments.

| Variant | Rheobase  (pA) | Threshold  (pA) | Width  (ms) | 90 to 10 % Decay  (ms) |
| --- | --- | --- | --- | --- |
| WT  n  P value | 6.40 ± 0.58  10  - | −33.27± 1.08  10  - | 1.70 ± 0.09  10  - | 3.04 ± 0.28  10  - |
| T162I  n  P value | 3.45 ± 0.28^***^  11  0.0004 | −46.12± 1.51^****^  11  <0.0001 | 1.38 ± 0.03^*^  11  0.041 | 2.27 ± 0.06^*^  11  0.018 |
| I236V  n  P value | 5.14 ± 0.73^NS^  7  0.308 | −37.19± 1.14^NS^  7  0.189 | 1.55 ± 0.05^NS^  7  0.661 | 2.72 ± 0.17^NS^  7  0.665 |
| P1345S  n  P value | 5.00 ± 0.44^NS^  6  0.259 | −35.31± 1.01^NS^  6  0.754 | 1.51 ± 0.07^NS^  6  0.504 | 2.46 ± 0.18^NS^  6  0.205 |
| R1636Q  n  P value | 3.33 ± 0.57^***^  9  0.0004 | −40.21± 1.68^**^  9  0.0028 | 2.69 ± 0.16^****^  9  <0.0001 | 5.01 ± 0.23^****^  9  <0.0001 |

Data are represented as mean ±SEM. In the AIS compartment model, the virtual Nav1.6 channel conductance (gNav1.6) and the virtual potassium channel conductance (gKv) values were set to gNa_v_1.6 = 0 and gK_v_ = 2. Firing was elicited by depolarizing step current stimuli in 2-pA increments. The first action potential elicited by a current step 2 pA above the rheobase was analysed. Statistically significant differences between the action potential characteristics of the wild-type (WT) and mutant channels were determined using one-way ANOVA followed by Dunnett’s post-hoc test (^*^P < 0.05, ^**^P < 0.01, ^***^P < 0.001, and ^****^P < 0.0001. Abbreviations: n, number of independent experiments; NS, statistically not significant difference compared to WT.

**Table A.3.** Selected biophysical characteristics of WT, T162I, I236V, P1345S, and R1636Q Na_v_1.1 channel variants in this study and reported previously.

| Na_v_1.1 channel  variant | This study | (Berecki et al., 2019) | (Brunklaus et al., 2022) | (Clatot et al., 2022) | (Matricardi et al., 2023) |
| --- | --- | --- | --- | --- | --- |
| WT  Expression system  Subunits expressed  I_density_ (pA/pF)  V_0.5,act_ (mV)  V_0.5 inact_ (mV)  I_NaP_ (% peak)  τ recovery^♣^ (ms) | CHO  α  535.2 ± 76  −17.6 ± 1.1  −49.7 ± 0.8  1.86 ± 0.2  1.08 ± 0.1 | CHO  α  530.5 ± 86  −18.5 ± 0.7  −48.8 ± 0.7  nr  1.39 ± 0.1 | HEK  α  194.51 ± 34  -25.29 ± 1.3  -60.8 ± 1.3  2.44 ± 0.9  ^Ψ^3.73 ± 0.2 | HEK  α + β_1_ + β_2_  90.0 ± 16  −21.2 ± 1.4  −61.3 ± 1.0  4.7 ± 0.9  nr | HEK  ^♠^α  153.6 ± 20  −27.6 ± 1.2  −63.3 ± 0.9  1.6 ± 0.3  ^Ψ^3.1 ± 0.3 |
| T162I  Expression system  Subunit transfected  I_density_ (pA/pF)  V_0.5,act_ (mV)  V_0.5 inact_ (mV)  I_NaP_ (% peak)  τ recovery^♣^ (ms) | CHO  α  880.4± 95  −24.0 ± 1.1*  −53.7 ± 1.1*  2.22 ± 0.2  1.34 ± 0.08 | ND  -  -  -  -  -  -  - | ND  -  -  -  -  -  -  - | ND  -  -  -  -  -  -  - | HEK  ^♠^α  147.0 ± 11  −34.2 ± 0.9*  −61.7 ± 1.1  2.2 ± 0.4  ^Ψ^2.1 ± 0.2 |
| I236V  Expression system  Subunits expressed  I_density_ (pA/pF)  V_0.5,act_ (mV)  V_0.5 inact_ (mV)  I_NaP_ (% peak)  τ recovery^♣^ (ms) | CHO  α  726.6 ± 147  −21.2 ± 1.1  −46.5 ± 1.5  3.56 ± 0.3*  1.04 ± 0.08 | ND  -  -  -  -  -  -  - | HEK  α  184.66 ± 37  -27.65 ± 1.3  -59.1 ± 1.7  4.23 ± 1.1*  ^Ψ^4.02 ± 0.5 | ND  -  -  -  -  -  -  - | ND  -  -  -  -  -  -  - |
| P1345S  Expression system  Subunits expressed  I_density_ (pA/pF)  V_0.5,act_ (mV)  V_0.5 inact_ (mV)  I_NaP_ (% peak)  τ recovery^♣^ (ms) | CHO  α  518.0 ± 132  −23.8 ± 1.1*  −46.7 ± 1.7  3.25 ± 0.3*  1.01 ± 0.07 | ND  -  -  -  -  -  -  - | ND  -  -  -  -  -  -  - | ND  -  -  -  -  -  -  - | ND  -  -  -  -  -  -  - |
| R1636Q  Expression system  Subunits expressed  I_density_ (pA/pF)  V_0.5,act_ (mV)  V_0.5 inact_ (mV)  I_NaP_ (% peak)  τ recovery^♣^ (ms) | CHO  α  544.1 ± 119  −24.9 ± 1.1*  −42.5 ± 1.1*  5.32 ± 0.5*  0.81 ± 0.04 | ND  -  -  -  -  -  -  - | HEK  α  101.80 ± 12  -26.51 ± 1.1  -60.7 ± 1.1  1.03 ± 0.2  ^Ψ^2.39 ± 0.3* | HEK  α + β1 + β2  126.9 ± 24  −27.2 ± 1.4*  ^Ω^−66.3 ± 1.1*  22.6 ± 4.0*  nr | ND  -  -  -  -  -  -  - |

Data are represented as mean ± SEM. *, asterisk indicate P < 0.05 versus control (WT); CHO, Chinese hamster ovary; §, cortical PV interneuron; HEK, human embryonic kidney (HEK-293T, also known as tsA-201); ^♠^, β_1_ and β_2_ subunit were co-expressed with α subunits and only tested in a subset of action potential clamp experiments; I_density_, current density; #, GABAergic interneuron model; I_NaP_, persistent sodium current; **^♣^**, ND, not determined; recovery from fast inactivation; nr, not reported; but ^Ψ^, recovery from fast inactivation was determined from a holding potential of −80 mV, which results in a slower time course compared with recovery from a more negative holding potential (Berecki et al., 2010); V_0.5,act_, membrane potential for half-maximal activation;V_0.5,inact_, membrane potential for half-maximal inactivation. ^Ω^, the hyperpolarized shift of the inactivation is mainly due to the decrease of the slope of the steady-state inactivation curve compared with the WT.

In CHO cells expressing Na_v_1.1 channel variants, the voltage dependence of activation and inactivation is more depolarized compared to data obtained in HEK cells. However, there are also notable differences in the activation and inactivation properties or the magnitude of persistent current of Na_v_1.1 variants among studies using HEK cells, suggesting that the expression system is not the sole cause for differences in the biophysical properties. Various factors, including the transfection system used, cell culture conditions, the amount and quality of transfected cDNA, the transfection method, and/or the composition of solutions used for electrophysiological recordings can affect the biophysical properties of the expressed channels. Lipofectamine 3000 Reagent (this study), CaPO_4_ (Brunklaus et al., 2022; Matricardi et al., 2023), or PolyFect Reagent (Clatot et al., 2022) was used for transfections; The composition of extracellular solutions was similar in all studies, whereas the intracellular solution included ATP (this study) or lacked ATP (Brunklaus et al., 2022; Clatot et al., 2022; Matricardi et al., 2023). Despite the apparent differences between the biophysical characteristics in these studies, both mammalian cell lines are considered as adequate for studying neuronal sodium channels.

**Table A.4.** General clinical features and treatment response of NDEEMA patients carrying the T162I mutation.

| Clinical features^§^ | Patient 1  This study | Patient 2  Brunklaus et al. 2022 | Patient 3*  Matricardi et al. 2023 |
| --- | --- | --- | --- |
| Inheritance/ Sex | De novo/ Female | De novo/ Male | De novo/ Female |
| Age at seizure onset | 2 d | 1 d | 1 d |
| Initial seizure type | T, AP, TC | T, AP | nr |
| Triggers | touch (esp. diaper change), not fever | Environ. stimuli,  Not fever | nr |
| Other seizure types | TC, ES (6 m), FC, focal motor SE | T, eyelid M, F, AtA, HC, SE, NCSE | F motor, F to bilateral, Tonic, Myoclonic, ES |
| Contractures, bone fractures | Cong. left hip dysplasia, cong. UL AGP | Cong. 4 limb AGP |  |
| Movement disorder (onset) | muscular hypotonia, peri-oral myoclonus, (6 months), transient nystagmus (6-12months), hyperkinesia/ chorea (18 m), | Hypertonia (birth), chorea, hyperkinesia (10 m) | Spastic quadriplegia |
| Age at last follow-up | 7y 7m | 2 y 6 m | 13 y 8 m |
| Development delay (onset) | Global delay (birth), motor regression (3 y), profound ID, NV, NA, | Global delay (birth), profound ID, NV, NA, no microcephaly | Severe ID |
| EEG | MFD, focal midline status epilepticus, GSW | MFD, migrating, GSW, NCSE | nr |
| MRI | normal (3d, 19m), delayed myelination (7m) | Normal 3 m & 6 m | nr |
| Seizure outcome | Intractable | Intractable |  |
| Seizure reduction | PB, TPM, LCM | OXC, PHT, TPM | CBZ |
| No effect | LEV, BRV, OXC, FEN, BR, KD | VPA, PHB, CLB, LTG, STP, FEN, ESM, KD | PB, CBZ, CLB, VGB, VPA, LEV, ESM, PER |
| Worsening | STP, CLB, CBD, RUF, LTG, AZA, CLH | No | nr |

NDEEMA, severe neonatal developmental and epileptic encephalopathy with movement disorder and arthrogryposis; §, list of clinical features according to Brunklaus et al (2022); *, also reported as a Dravet syndrome patient (Cetica et al., 2017); d, day; m, month; y, year; AP: apnoea, AtA, atypical absence; Cong., congenital; ES. Epileptic spasms; F, focal; M, myoclonus; NCSE, non-convulsive status; SE, status epilepticus; T, tonic; TC, tonic-clonic; AGP, FC, focal-clonic; arthrogryposis; UL, upper limb; ID, intellectual disability; NV, non-verbal; NA, non-ambulant; GSW, generalised spike-wave, MFD, multi-focal discharges; AZA, acetazolamide; BR, Potassiumbromide; BRV, brivaracetam; CBD, Cannabidiol; CBZ, carbamazepine; CLH, chloral hydrate; CLB, clobazam; ESM, ethosuximide; FEN, fenfluramine; KD, ketogenic diet; LCM, lacosamide; LEV, levetiracetam; LTG, lamotrigine; OXC, oxcarbazepine; PER, perampanel; PB, phenobarbital; RUF, rufinamide; STP, stiripentol; TPM, topiramate; VPA, valproate; VGB, vigabatrin; N/A: not applicable; nr, not reported.
